# Distant Site Mutations in Clinical TEM Beta-Lactamase Variants Enhance Non-Covalent Binding to Ceftazidime: Insights from Spectroscopic and Biophysical Investigations

**DOI:** 10.1101/2025.08.06.661929

**Authors:** Sandip Kumar Mukherjee, Padmaja Prasad Mishra, Mandira Mukherjee

## Abstract

β-lactamases retain the central armamentarium against the β-lactams, resulting in surge of antibiotic resistance, primarily due to the hydrolysis of the amide bond of the four-membered ring. This study aimed to investigate the binding interactions of ceftazidime (CAZ) to TEM β- lactamase variants with distant site mutations isolated from clinical setting to explore the cause of their selection and dissemination due to indiscriminate use of β-lactams. Absorbance and fluorescence spectroscopy along with biophysical experimentation indicated facilitated binding of CAZ to the active site of the mutants than the wild type. The CAZ-TEM β-lactamase mutant interactions were predominantly hydrophobic compared to H-bonding and van der Waals forces in the CAZ-wild type complex. Additionally structural alteration to justify more rigid binding of CAZ to the mutants in contrast to the wild type enzyme was established *in silico*. Therefore, acquisition of distant site mutations with respect to the active site of the β-lactamase variants that rendered conformational flexibility to accommodate CAZ was evidenced. This study provided an insight to the bioactive interaction of CAZ with TEM β-lactamase variants that probably facilitated their selection from clinical settings in response to rampant usage of β-lactams.

## 1. Introduction

Persistence of bacterial resistance against antimicrobial agents is a global health threat (Avci et al. <u>201</u>6). Production of active site serine β-lactamase enzymes is the major mechanism of antibiotic resistant in uropathogenic *Escherichia coli* (UPEC) that catalyze the hydrolysis of β- lactam antibiotics. (Chinemerem et al. 2022,Natalia Díaz et al.2003,Stojanoski et al.2015). Currently, based on Ambler classification scheme, β-lactamases can be grouped into four classes A, B, C and D. Class A, C, and D are serine β-lactamases (non-metallo) that utilize the serine residue to catalyze the hydrolysis of antibiotics, whereas class B enzymes are metallo-β- lactamases and need zinc ions in the active site to attack the β-lactam ring(Stojanoski et al.2015).In the case of Class A β-lactamases which is prime concerned for the clinicians due to its wide range of hydrolyzing activity against antibiotics mostly against β-lactam antibiotics where activation of catalytic Ser70 conserved residue initiates nucleophilic attack on the β- lactam carbonyl carbon and opening of the β-lactam ring to form a covalently linked acyl-enzyme intermediate and ultimately the attack leads to deacylation of the acyl-enzyme species resulting inactivation of antibiotic and regeneration of the free enzyme (Stojanoski et al.2015).TEM1 β-lactamase was identified as one of the most evolved Class A β-lactamases in hospital infection, mediated by plasmid found in *E. coli* resistant to antibiotics and also considered as the precursor of the TEM enzyme family with three major types of activity; the broad-spectrum β-lactamases, the extended spectrum β-lactamases (ESBLs) and the inhibitor resistant β-lactamases. The emergence of resistance mediated by this TEM enzyme family were most explored β-lactamases till date and it was found that their extensive structural adjustment by their sustainable mutational modifications led to the generation of distinct multiple variants with in-depth structural diversity (Singhet al.2012). To overcome β-lactam resistance extensive studies were carried out against the acylation mechanism and two types of β-lactams were introduced such as extended spectrum cephalosporins; and β-lactamase inhibitors, such as clavulanic acid, sulbactam and tazobactam [(Kumar et al.2014). Furthermore, the structural alterations in Class A β-lactamases impair the antibacterial activities of the inhibitors by covalent interaction between the active site residues in the former with the inhibitory molecules resulting in resistance against them (Drawz and Bonomo 2010). Till date ∼200 TEM-variants have been reported that confers resistance to novel β-lactams and their inhibitor combinations which were introduced in the past three decades (Barlow et al, 2010. Pimenta et al,2014). Studies from the different part of the world showed that the indiscriminate use of β-lactam-β-lactamase inhibitor combination has increased the number of TEM β-lactamase variants; the extended-spectrum (ES) and inhibitor-resistant phenotypes (IR) (Tooke et al.2019,Cantón et al. 2008). Previous reports showed that mutation or substitution in the active site Ω-loop region of TEM β-lactamase not only increased the resistance phenotype against cephalosporins but also increased the structural flexibility imparting enlargement of the active site pocket loop that can accommodate newer β- lactams irrespective of their -R group thus decreasing the activity of oxyimino-cephalosporin, ceftazidime (CAZ), and all other β-lactam-β-lactamase inhibitors (Natalia Díaz et al.2003,Stojanoski et al.2015). Fluorescence techniques were widely used as one of the important biophysical application that paved sensitivity, reliability and rapidity in the field of drug-protein interactions study (Moller et al.2002,Kumar et al. 2024).

There are not many Indian studies that described existence and characterization of TEM β- lactamase mutants that harbor mutations at sites other than the active site of the enzyme isolated from clinical origin as a result of random drug administration. An earlier study from our laboratory, demonstrated identification and characterization of three novel TEM β-lactamase mutants; pUE184TEM (M184), pUE203TEM (M203), pUE210TEM (M210) from clinical uropathogenic *E. coli* isolates with mutation at residues other than those that constituted the active site of the enzymes (Mukherjee S K et al, 2018). In the present study bioactive interactions of CAZ with the mutant β-lactamases under physiological conditions was investigated. The role of distant site mutations on the conformational flexibility in the mutants to accommodate CAZ was explored by using a collection of biophysical and *in silico* methods to obtain an insight into selection and dissemination of the resistant β-lactamases in response to uncontrolled β-lactam administration.

## 2. Materials and Methods

### Absorbance Spectroscopy

Absorbance spectral measurement wasrecorded in UV–vis Spectrometer at a scan rate of 480 nm/min with 1 cm-path-lengthquartz cuvette in the visible wavelength range of 200-800nmat different temperatures (298K, 303K, 308K, 311K).The titrations of the experiment were conducted on 5 μM wild type TEM β-lactamase and its variants (M184, M203, M210) (Mukherjee S K et al, 2018) in presence ofascendingconcentration of CAZ (5 μM-20μM) at respective temperatures.The inner filter effect was eliminated using the relationship; I_cor_ = I_obs_e^(Aex+Aem)/2^,where the corrected fluorescence intensity is denoted as I_corr_, and observed fluorescence intensity is denoted as I_obs_. The A_ex_ and A_em_ denote absorbance values at excitation and emission wavelengths, respectively (Swain et al. 2020)

### Fluorescence Spectroscopy

Fluorescence emission spectrum was recorded in the range of280–600 nm using Hitachi F-7000 Spectrofluorimeter with excitation at 295 nm. Both the excitation and emission bandwidth were fixed at5 nm. Quenching experiment was carried out in presence of increasing concentration of CAZ (5 μM-20μM) solution at different temperatures (298K, 303K, 308K, 311K) to 5 μM concentration of the wild type and the mutant β-lactamases.Static fluorescence quenching was determined by using the Stern–Volmer (SV) equation from steady state measurements F_0_/F = 1 + K_sv_(Q), where F_0_ and F denote the steady-state fluorescence emission intensities in the absence and in presence of CAZ. K_SV_ is the Stern–Volmer constant and (Q) is the concentration of the quencher. Collisional or dynamic fluorescence quenching was determined using the modified Stern-Volmer equation; log ((F_0_ -F)/F)) = logK_a_ + n log (Q), where K_a_ denoted the binding constant and n the number of binding sites.

### Fluorescence lifetime measurements

The fluorescence lifetime(τ) measurements determine energy transfer at the molecular level between interacting species. Time correlated single photon counting (TCSPC) was used to measure the fluorescence life times of wild type and mutant β-lactamases in gradually increasing concentration of CAZ (0 μM-20μM) in sodium phosphate buffer (pH 7.4) at 298K (Maldal S K et al, 2012).

### Quantum yield and fluorescence anisotropy

The quantum yield (Φ) or quantum efficiency of CAZ at 5µM concentration in sodium phosphate buffer (pH 7.4) was recorded by monitoring the fluorescence emission intensityin presence of increasing concentrations of the wild type and mutant β lactamases (0-20 µM)at 298K (Mishra et al, 2006). The fluorescence anisotropy (r) measurement of CAZ at 5μM concentration in 5mM sodium phosphate buffer (pH 7.4) was recorded against addition of ascending concentration of wild type and mutant proteins (5μM-20μM) in Fluromax Spectrofluorometer. All measurements were conducted at 298K at 3 minutes of interval.

### Thermodynamic properties

Van’t Hoff isotherm equation was used to determine the thermodynamic parameter (kinetic and equilibrium properties) that govern the interaction between the ligands (CAZ) and the proteins (TEM and its mutants). The values of ΔH^0^ and ΔS^0^ were obtained from the *lnK* vs. 1/T plot, where ΔG^0^, ΔH^0^, and ΔS^0^ are depicted as standard enthalpy change, standard free energy change, and standard entropy change respectively.

### Circular Dichroism (CD) Spectroscopy

Far and Near UV-CD spectra was recorded at room temperature for β-lactamases (TEM and its mutants) in presence of ascending concentration of ceftazidime ranges from 0 μM-20μM solutions using JASCO-J815 spectrometer with a temperature controller at 298K in a quartz cuvette of 1 mm path length. Each final spectrum was recorded as an average over three scans. The far and near-UV-CD region was recorded as 250–360 nm and 360 nm to 500 nm.CD results were expressed as mean residue elasticity (MRE) in degree cm^2^ mol^-1^. The secondary structures (α-helix, β-sheet, random coils) were calculated from respective MRE values (Bertucci et al.2010) **Raman spectroscopy**

Raman scattering was recorded for β-lactamases (TEM and its mutant variants) in presence of ascending concentration of ceftazidime ranges from 0μM-20μM. The spectral range was taken for500–4000 cm^−1^ with an integration time of 20 s using a 670 nm red laser excitation, at 10 mW power (Miani et al.2007).

### Förster resonance energy transfer (FRET) computation

FRET involves a distance dependent physical method of non-radiative transfer of energy between an excited fluorophore (donor) and another fluorophore (acceptor) by intermolecular dipole-dipole interactions. The energy transfer efficiency (E) signified the fraction of photons absorbed by the donor moiety which was transferred to the acceptor. E= 1-(F/F_0_) = R_0_^6^/(R_0_^6^ + r^6^), where, r represented the distance between the donor and acceptor and R_0_ represented the critical energy transfer distance, at which 50% of the excitation energy was transmitted from the former to the latter. R_0_ was evaluated from the following equation; R_0_^6^ = 8.79 x 10^-25^K^2^N^-4^, where K^2^ was the spatial orientation factor of the altered dipoles of the acceptor and the donor, N represented the refractive index of the medium, denoted the fluorescence quantum yield of the donor in absence of the acceptor. (Chakraborty et al.2015).

### MD simulation

Molecular dynamics and simulation was performed on the TEM β-lactamase (1ZG4) and its variants (Mukherjee S K et al, 2018) in presence of CAZ to determine the binding stability of the protein–drug docked complex using Desmond v3.6 Package. Topology of the protein molecules were generated by optimized potentials for liquid simulation (OPLS) force field. Simple point charge (SPC) water model was used to perform the protein–drug complexes Simulation with a distance of 10 Å between the complex and the periodic box. System was neutralized by adding Na+/Cl− ions and placed randomly in the solvated system. To minimize and relax the protein/protein−drug complex under NPT ensemble, the default protocol of Desmond was used followed by total of nine stages among which two minimization and four short simulation (equilibration phase) steps was involved before starting the actual production time. System temperature and pressure was 300 K and 1 atmosphere, respectively, using Nose−Hoover temperature coupling and isotropic scaling, and the simulation was performed for 100 ns NPT production simulation and saving the configurations at 10 ns intervals (Mukherjee S K et al, 2021).

### Simulation trajectory analysis

Simulation trajectory files were analyzed using Desmond module programs i.e. simulation quality analysis (SQA), simulation event analysis (SEA), and simulation interaction diagram (SID) to calculate energies, root-mean-square deviation (RMSD), root-mean-square fluctuation (RMSF), the radius of gyration (Rg) and the secondary structure elements (SSEs) which pave the protein structure stability. SQA was used to qualitatively validate the system stability throughout the simulated length of chemical time for the given temperature, pressure, and volume of the total simulation box, SEA was used to analyze each frame of the simulated trajectory output, whereas total SSE change in the protein structure in presence of the drug during the simulation time was estimated using SID (Abdul et al, 2016, (Matyus et al, 2006)).

## Results

### Spectroscopic measurements

UV-Vis absorption spectra of wild type and mutant β-lactamases (**Fig. 1A-D**) were recorded in presence of increasing concentrations of ceftazidime, at 298K. All protein showed a broad band at 280 nm while ceftazidime had a strong absorbance at 267 nm and a moderate absorbance at 310 nm. With gradual increase of CAZ concentration, the absorbance spectra of wild β-lactamase showed enhancement of absorption intensity.

**Fig. 1.**
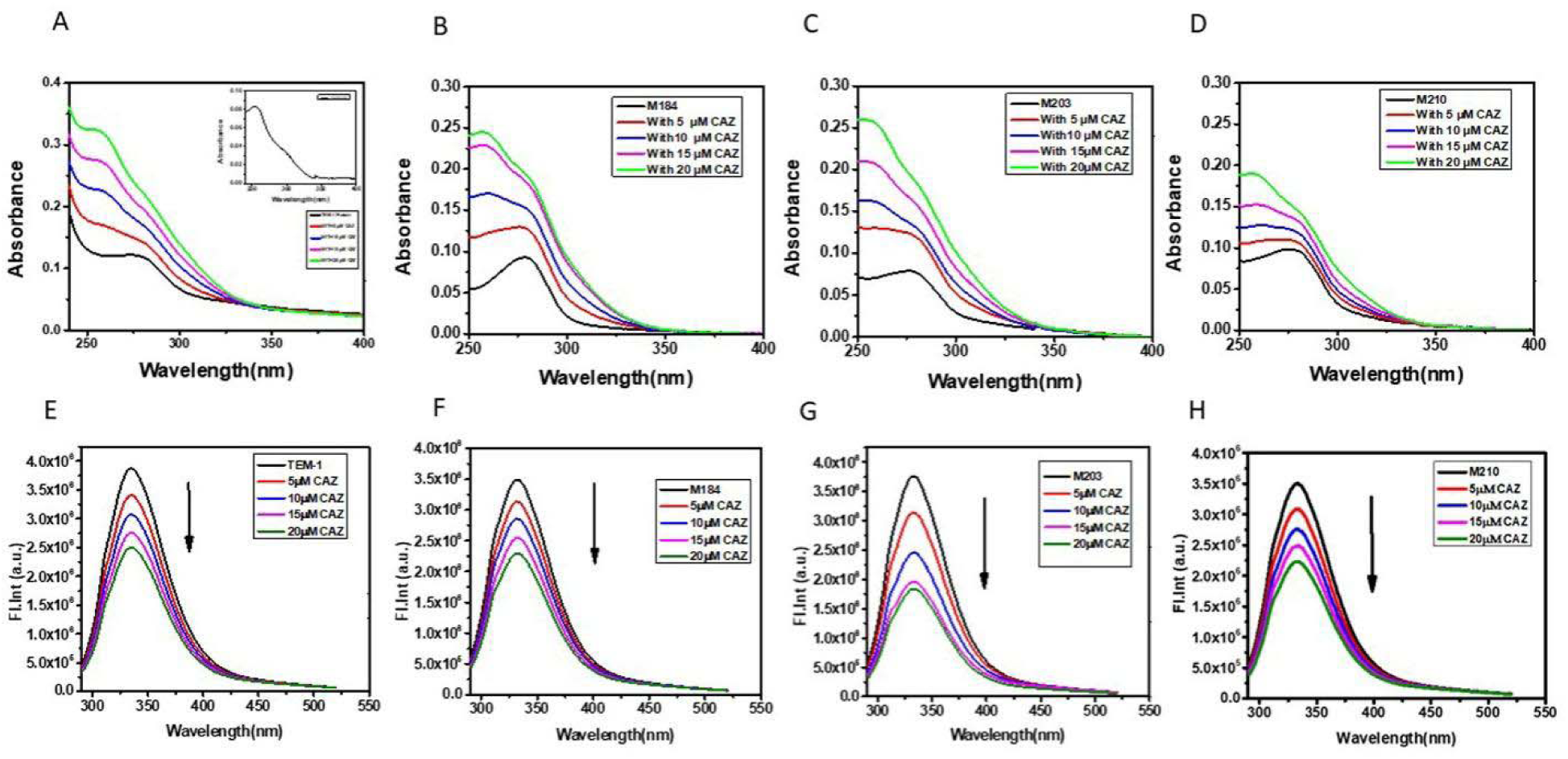
Steady-state UV–vis absorption spectra and fluorescence emission spectra of TEM-β-lactamase (A,E) and mutant β-lactamases 184M (B,F), M203 (C,G), M210 (D,H) at 5 µM concentration of the respective protein in solution in presence of increasing concentration of CAZ (0-20 µM). Inset in (A) represented steady-state UV–vis absorption spectra of CAZ at 20 µM at 298 K.

The possibility of ground state complex formation couldnot be ruled out from the above absorption spectra. Additionally, from **Fig. S1** it was observed that the spectra of the mixture of CAZ and wild type β-lactamase (**curve 5**) was much intense than intensity of spectra corresponding to the summationof the two absorption bands of the individual reacting components (**curve 3**). The **curve 4** obtained by subtracting the **curve 3** from the **curve 5** indicated formation of ground state complex between the two species.

The UV-Vis absorption spectra of the wild type and mutant β lactamases in presence of CAZ were recorded at 303 K, 308 K and 311 K respectively which indicated that the proteins-CAZ complexes were thermally stable (**Fig. S2**).

The fluorescence spectra of wild type and mutant β-lactamases at pH 7.5in the presence of 5, 10, 15, 20μM CAZ at 298K showed strong fluorescence intensity at 340 nm upon excitation at 280 nm which gradually decreased upon addition of CAZ with a slight red shift at 298 K (**Fig. 1E-H, Fig. S3**). Similar comparable fluorescence spectra of the β-lactamases at the different temperatures (303K.308K, 311K) were also recorded (data not shown).

The quenching phenomenon exhibited in the fluorescence emission spectra of the proteins and protein –CAZ complexes were analyzed by Stern–Volmer (SV) plot. The non-linearity observed in the SV plot at 298K, 303K, 308K, 311K (**Fig. 2A-D**) indicated a combination of static (ground state) and dynamic (excited state) quenching which was validated by modified SV plots (**Fig. 2E-H**). Additionally, SV plots of quenching of the wild type in presence of increased concentration of CAZ at different temperatures indicated a gradual shift towards dynamic mode although mutants exhibited a decrease in their dynamic mode of quenching.

**Fig.2.**
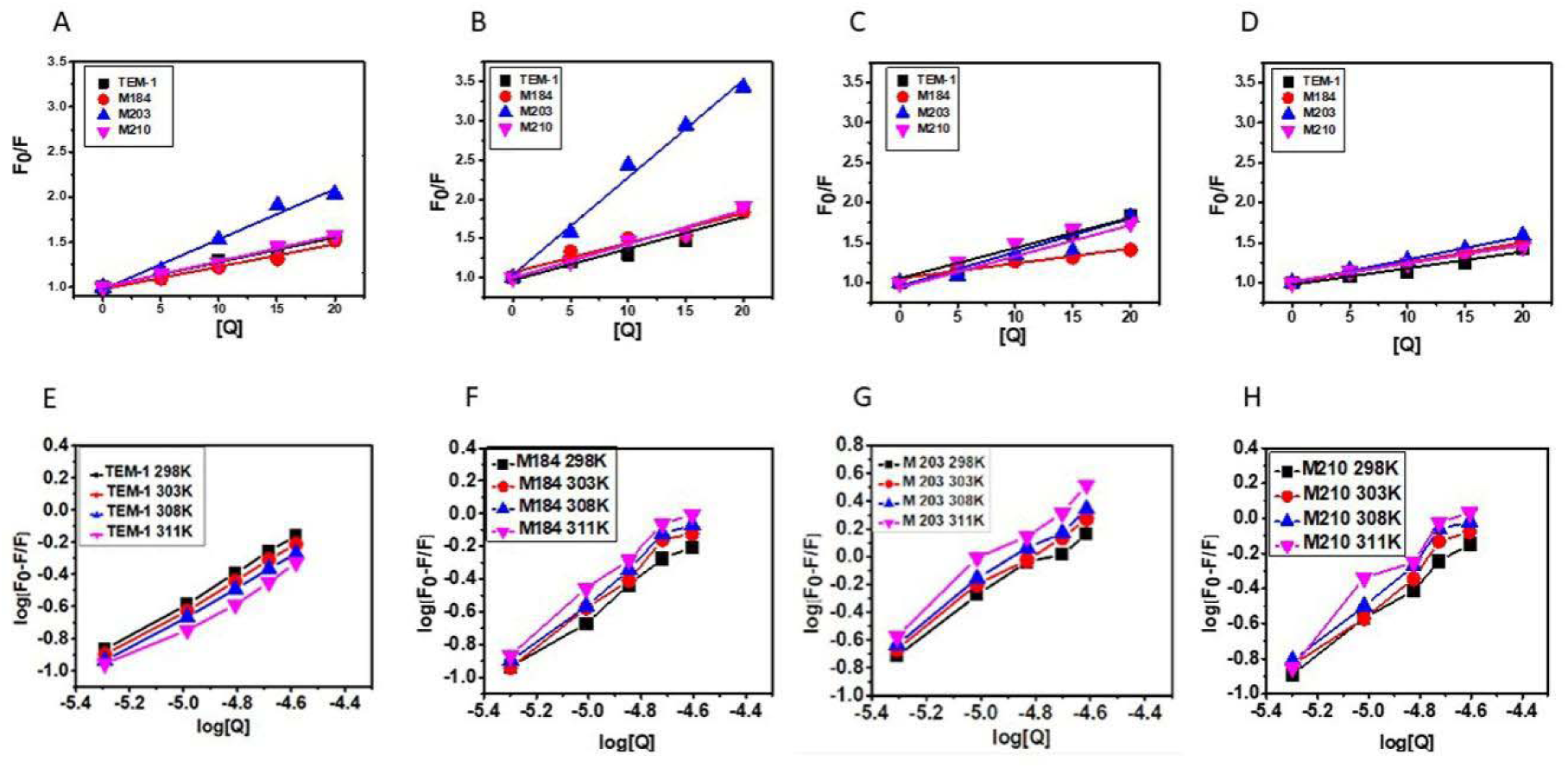
Stern-Volmer plot and Modified Stern-Volmer plot of Fluorescence quenching of TEM-β lactamase (wild type protein) and mutant β lactamases (M184, M203, M210) at 5 µM concentration in the presence of CAZ (0-20 µM) at 298 K (A,E), 303K (B,F), 308K (C,G) and 313K (D,H). Nonlinear fitting analysis was performed by plotting F0/F against [Q] and Log F0-F/F1 against Log{Q} respectively.

**Fig.3.**
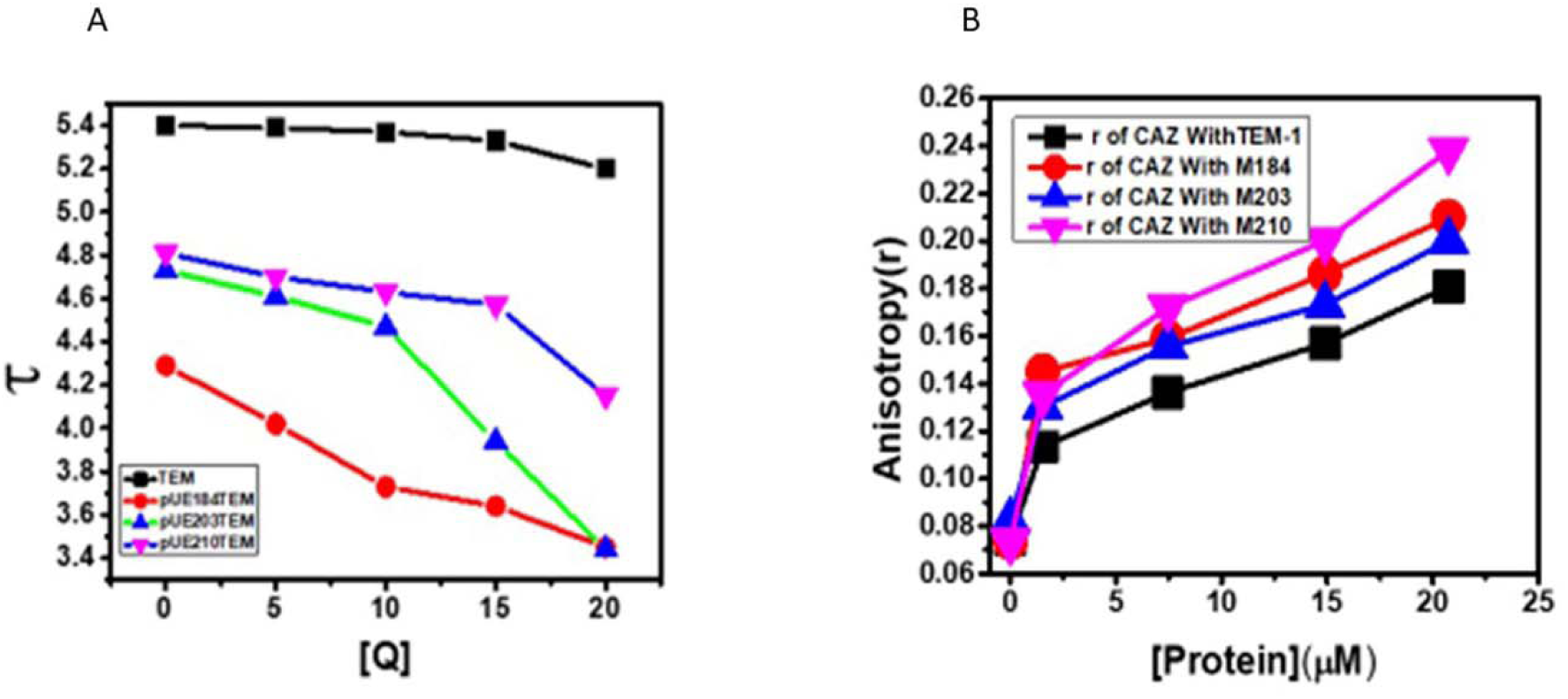
Fluorescence lifetime (τ) of TEM-β lactamase (wild type protein) in comparison with mutant β lactamases (M184, M203 and M210) at 5 µM concentration in the presence of CAZ (0-20 µM) at temperature 298K (A) and Fluorescence anisotropy (r) plot of CAZ at 5 µM concentration in presence of increasing concentrations (5-20 µM) of the β lactamases (TEM 1, M184, M203, M210) (B).

Furthermore, fluorescence lifetime changes of the wild type and the mutant proteins in presence of different concentrations of CAZ also exhibited a steady decrease (**Fig. 3A**, **Table 1**).

**Table 1:**
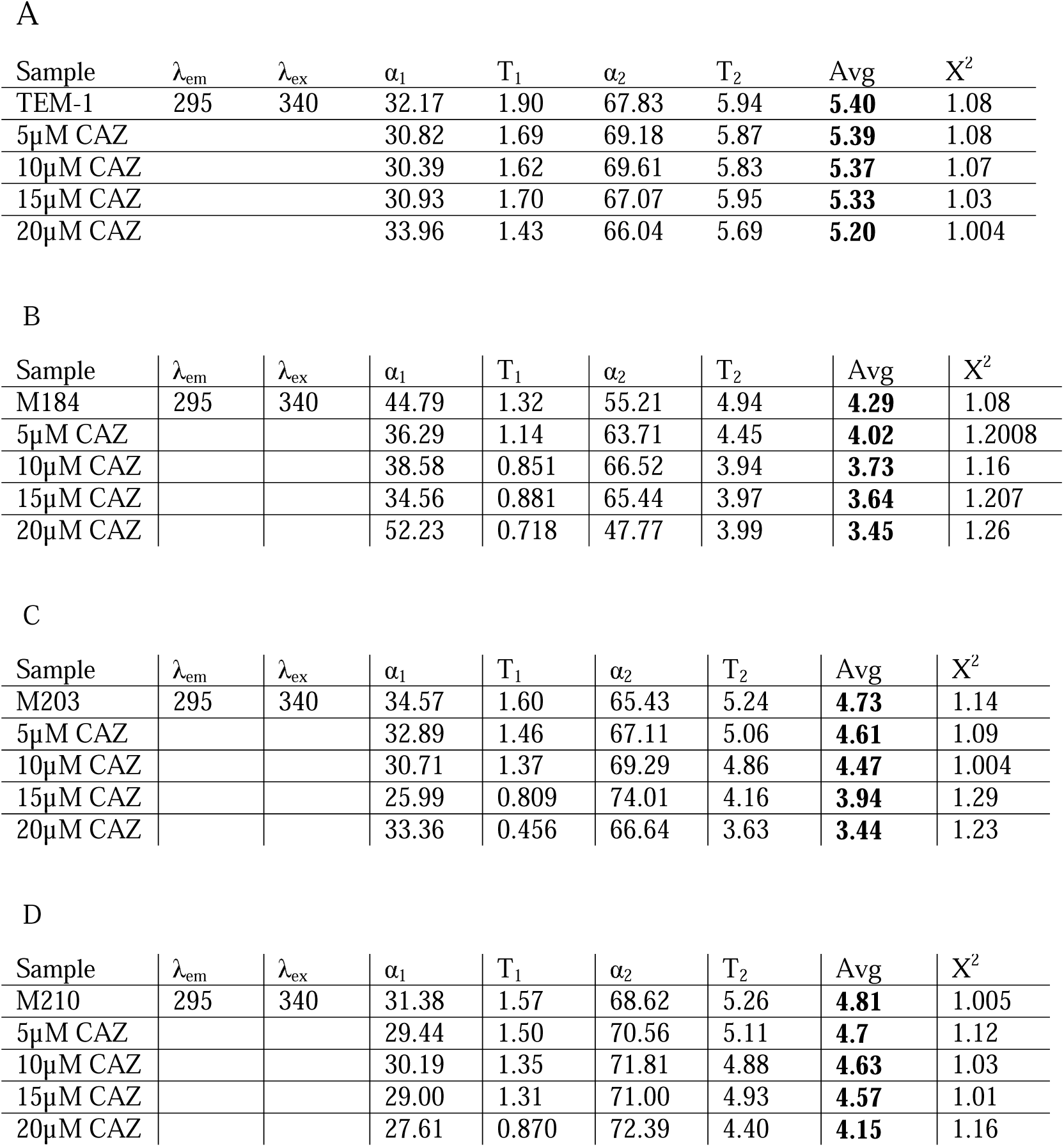
TCSPC life time fluorescence decay profile of (A) TEM-1(wild type protein) and mutant β-lactamases (B) M184, (C) M203 and (D) M210 at 0.5 mM concentration in the absence and presence of Ceftazidime (CAZ).

| Sample | $\lambda_{em}$ | $\lambda_{ex}$ | $\alpha_1$ | $T_1$ | $\alpha_2$ | $T_2$ | Avg | $X^2$ |
| --- | --- | --- | --- | --- | --- | --- | --- | --- |
| TEM-1 | 295 | 340 | 32.17 | 1.90 | 67.83 | 5.94 | **5.40** | 1.08 |
| 5μM CAZ | | | 30.82 | 1.69 | 69.18 | 5.87 | **5.39** | 1.08 |
| 10μM CAZ | | | 30.39 | 1.62 | 69.61 | 5.83 | **5.37** | 1.07 |
| 15μM CAZ | | | 30.93 | 1.70 | 67.07 | 5.95 | **5.33** | 1.03 |
| 20μM CAZ | | | 33.96 | 1.43 | 66.04 | 5.69 | **5.20** | 1.004 |

B
| Sample | $\lambda_{em}$ | $\lambda_{ex}$ | $\alpha_1$ | $T_1$ | $\alpha_2$ | $T_2$ | Avg | $X^2$ |
| --- | --- | --- | --- | --- | --- | --- | --- | --- |
| M184 | 295 | 340 | 44.79 | 1.32 | 55.21 | 4.94 | **4.29** | 1.08 |
| 5μM CAZ | | | 36.29 | 1.14 | 63.71 | 4.45 | **4.02** | 1.2008 |
| 10μM CAZ | | | 38.58 | 0.851 | 66.52 | 3.94 | **3.73** | 1.16 |
| 15μM CAZ | | | 34.56 | 0.881 | 65.44 | 3.97 | **3.64** | 1.207 |
| 20μM CAZ | | | 52.23 | 0.718 | 47.77 | 3.99 | **3.45** | 1.26 |

C
| Sample | $\lambda_{em}$ | $\lambda_{ex}$ | $\alpha_1$ | $T_1$ | $\alpha_2$ | $T_2$ | Avg | $X^2$ |
| --- | --- | --- | --- | --- | --- | --- | --- | --- |
| M203 | 295 | 340 | 34.57 | 1.60 | 65.43 | 5.24 | **4.73** | 1.14 |
| 5μM CAZ | | | 32.89 | 1.46 | 67.11 | 5.06 | **4.61** | 1.09 |
| 10μM CAZ | | | 30.71 | 1.37 | 69.29 | 4.86 | **4.47** | 1.004 |
| 15μM CAZ | | | 25.99 | 0.809 | 74.01 | 4.16 | **3.94** | 1.29 |
| 20μM CAZ | | | 33.36 | 0.456 | 66.64 | 3.63 | **3.44** | 1.23 |

D
| Sample | $\lambda_{em}$ | $\lambda_{ex}$ | $\alpha_1$ | $T_1$ | $\alpha_2$ | $T_2$ | Avg | $X^2$ |
| --- | --- | --- | --- | --- | --- | --- | --- | --- |
| M210 | 295 | 340 | 31.38 | 1.57 | 68.62 | 5.26 | **4.81** | 1.005 |
| 5μM CAZ | | | 29.44 | 1.50 | 70.56 | 5.11 | **4.7** | 1.12 |
| 10μM CAZ | | | 30.19 | 1.35 | 71.81 | 4.88 | **4.63** | 1.03 |
| 15μM CAZ | | | 29.00 | 1.31 | 71.00 | 4.93 | **4.57** | 1.01 |
| 20μM CAZ | | | 27.61 | 0.870 | 72.39 | 4.40 | **4.15** | 1.16 |

### Fluorescence quantum yield and fluorescence anisotropyof CAZ

Absolute fluorescence quantum yield was also analyzed for small molecule i.e. CAZ to measure changes in radiative rates from changes in nonradiative rates occurring in terms of interaction. The quantum yields of the CAZ molecule increased with the increase in concentration of proteins (wild and all three mutant β-lactamases). The quantum yields of CAZ increased gradually from 0.08 to 0.27, 0.36, 0.36 and 0.37 for the wild type, M184, M203 and M210 β-lactamase respectively (**Table 2**).Fluorescence anisotropy was also performed to measure the microenvironment around CAZ in terms of rotational diffusion and interactions. The anisotropy of the CAZ molecule was very low (r = 0.07). With increased concentration of the proteins (wild type and all three mutant β-lactamases), fluorescence anisotropy of CAZ increased gradually. At 20µM concentration of CAZ a steady increase in r value from 0.18, 0.20, 0.21 to 0.24 in presence of the respective β-lactamases (wild type, M184, M203, M210) respectively (**Fig. 3B**)

**Table 2:**
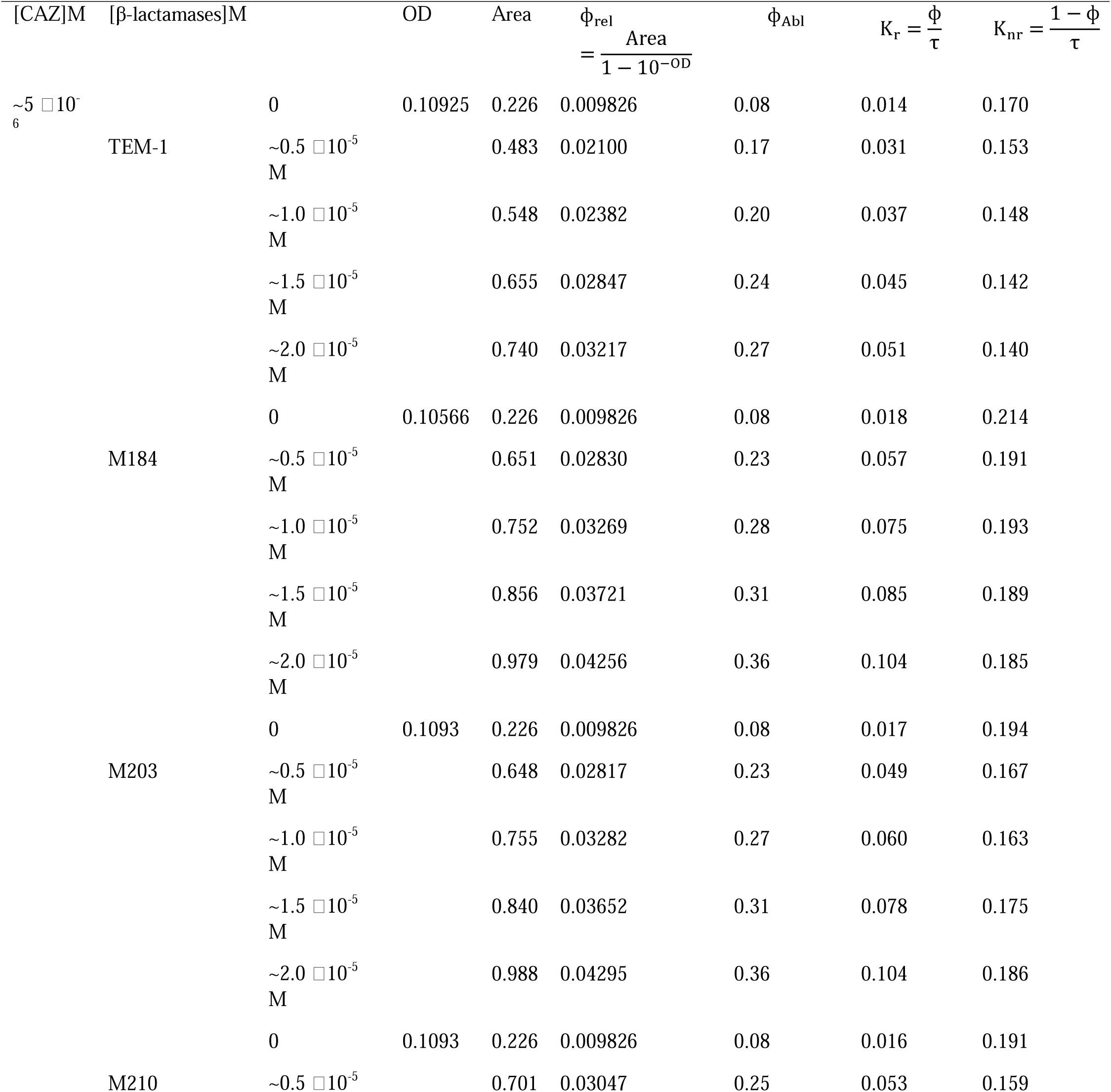
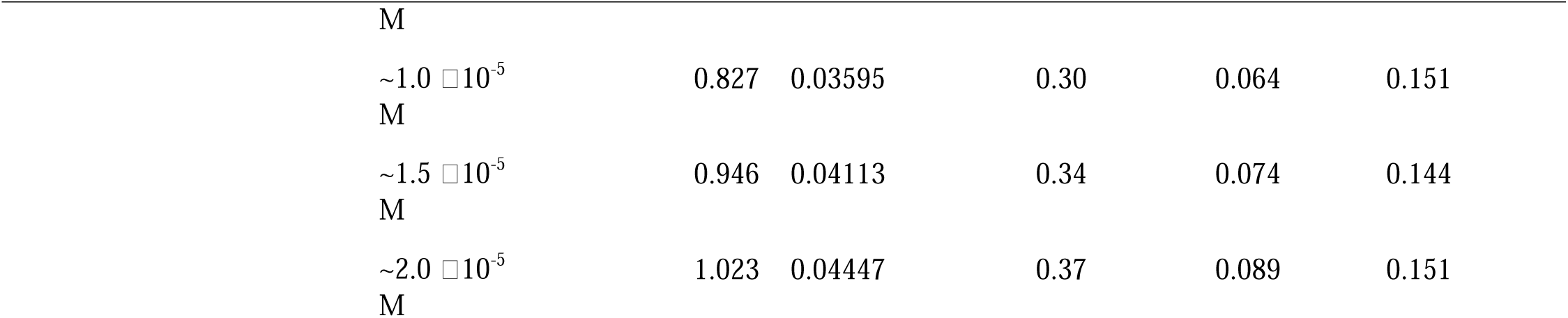
Quantum yield profile of TEM-1(wild type protein) and mutant β-lactamases (M184, M203 andM210)at different concentration in the absence and presence of Ceftazidime (CAZ).

### Binding constants, number of binding sites and mode of binding

The steady state fluorescence data were further evaluated by the binding affinity (K_b_) of CAZ molecule to the equivalent sites of wild type and mutant β-lactamases independently. For mutant β-lactamases a progressive decrease of binding affinity (K_b_) was observed with the increase of temperature whereas in case of wild type a vice versa affinity was observed in pH 7.5.(**Fig. 4**).

**Fig. 4.**
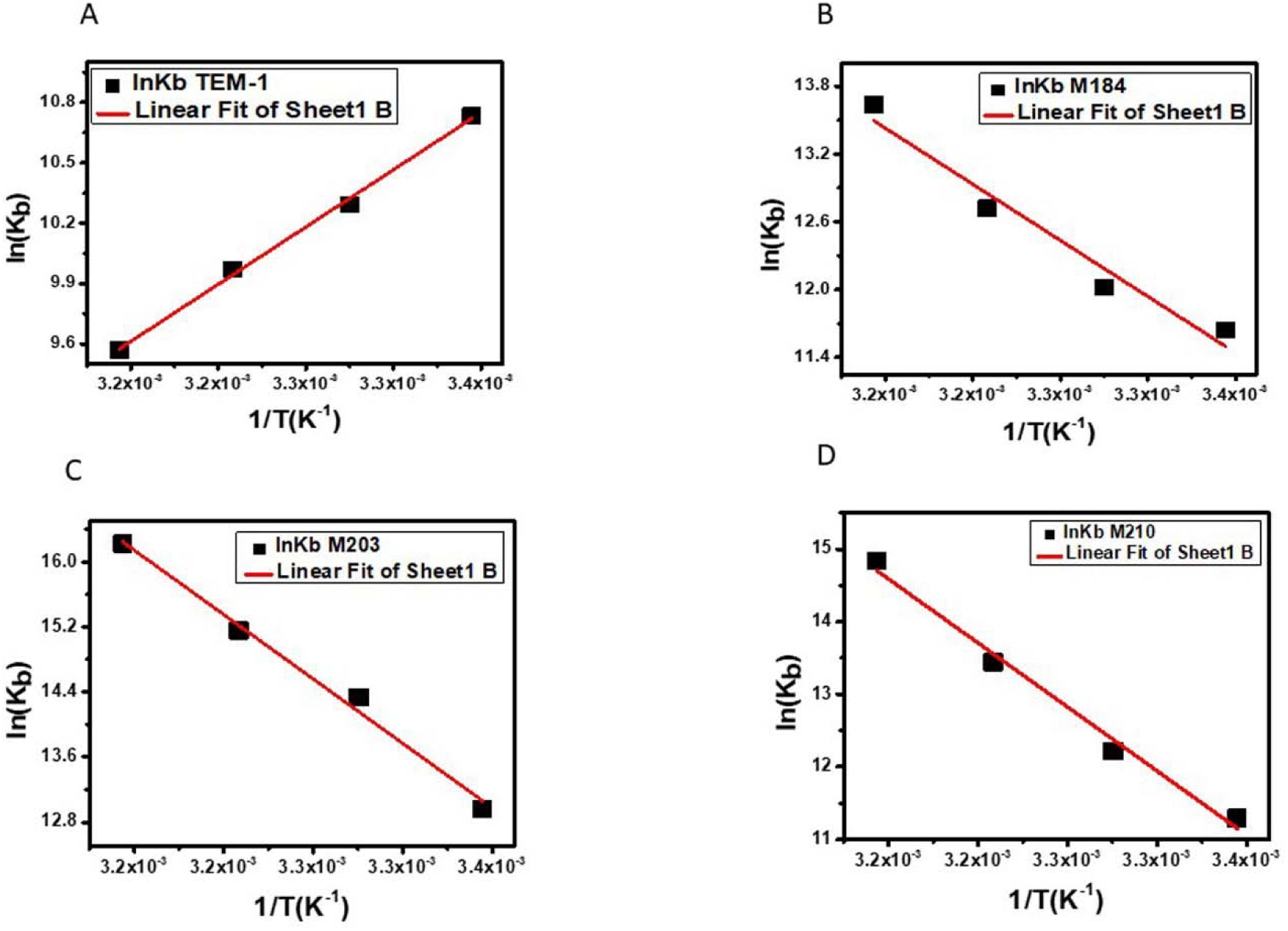
Van’t Hoff plot for the binding of TEM-β lactamases (wild type protein) (A) in comparison with mutant β lactamases M184(B), M203(C), M210(D) at 5 µM concentration in the presence of CAZ (0-20µM) concentration at different temperatures 298K, 303K, 308K and 311K.

The equilibrium interaction (such as, hydrogen bonds, van der Waals forces, electrostatic forces, and/or hydrophobic associations), between ligand (CAZ) and protein molecule were determined from the value of the thermodynamic parameter (ΔG^0^, ΔH^0^, ΔS^0^)using van’t Hoff isotherm equation from the lnK vs. 1/T plot. Significant negative ΔG^0^value, ΔH^0^< 0 and ΔS^0^> 0 indicated spontaneous interaction between the reacting components. Increase in negative ΔG^0^ value with increase in positive ΔH^0^ and ΔS^0^ values in the mutantβ-lactamases indicated discrete predominance of hydrophobic interaction in contrary to the wild type β-lactamase that exhibited van der Waals and H-bond interaction with high negative ΔH^0^and ΔS^0^ values compared to less negative ΔG^0^ value (**Fig. 4**, **Table 3**).

**Table 3:** Binding Parameters and Thermodynamic Parameters for the Binding of TEM-1(wild type protein) and mutant β-lactamases (M184, M203 and M210) with Ceftazidime (CAZ) at Different temperature.

| PROTEIN | Temperature | $\Delta G^0(\text{kJ mol}^{-1}\text{K}^{-1})$ | $\Delta H^0(\text{kJ mol}^{-1})$ | $\Delta S^0(\text{J mol}^{-1}\text{K}^{-1})$ | Pattern of Interaction |
| --- | --- | --- | --- | --- | --- |
| TEM-1 | 298K | -265.59378 | -591.644238 | -10.9412905 | V .W. Interaction and H-bond |
|  | 303K | -260.12313 |  |  |  |
|  | 308K | -254.652491 |  |  |  |
|  | 311K | -251.370103 |  |  |  |
| M184 | 298K | -284.81196 | 103.344092 | 44.2366739 | Hydrophobic |
|  | 303K | -306.93029 |  |  |  |
|  | 308K | -329.04863 |  |  |  |
|  | 311K | -342.31963 |  |  |  |
| M203 | 298K | -323.56295 | 164.577702 | 66.0852341 | Hydrophobic |
|  | 303K | -356.60557 |  |  |  |
|  | 308K | -389.64819 |  |  |  |
|  | 311K | -409.47376 |  |  |  |
| M210 | 298K | -276.32427 | 183.413275 | 70.8207056 | Hydrophobic |
|  | 303K | -311.73462 |  |  |  |
|  | 308K | -347.14498 |  |  |  |
|  | 311K | -368.39119 |  |  |  |

### Circular Dichroism (CD) Spectroscopy

Conformational changes of wild and mutant β-lactamases were further measured by CD spectroscopy in the absence and presence of CAZ at pH 7.5. Results showed that the mean residue ellipticity (MRE) of pure α-helix percentage of wild and mutant β-lactamases were decreased upon binding of CAZ demonstrated secondary structure alternations of the proteins in far-UV-CD region (185–250 nm) with two characteristic dips at 209 nm and 222 nm for all the proteins respectively. Moreover, in the near-UV-CD region spectra (240–320 nm) of the wild type and mutant proteins showed comparable minor alterations of their tertiary structure (**Fig. S4, Table 4**)

**Table 4:** Variation in a-helix% of TEM-1(wild type protein) and mutant β-lactamases (M184, M203 and M210) with increase concentration of Ceftazidime (CAZ).

| [β-lactamases]M | [CAZ]M | $\alpha$ – helix (%) |
| --- | --- | --- |
| TEM-1($\sim 5 \times 10^{-6}$ M) | 0 | 59.24 |
| | $\sim 0.5 \times 10^{-5}$ M | 41 |
| | $\sim 1.0 \times 10^{-5}$ M | 27.17 |
| | $\sim 1.5 \times 10^{-5}$ M | 26.72 |
| | $\sim 2.0 \times 10^{-5}$ M | 20.72 |
| M184($\sim 5 \times 10^{-6}$ M) | 0 | 44.10 |
| | $\sim 0.5 \times 10^{-5}$ M | 42.55 |
| | $\sim 1.0 \times 10^{-5}$ M | 34.17 |
| | $\sim 1.5 \times 10^{-5}$ M | 33.27 |
| | $\sim 2.0 \times 10^{-5}$ M | 14.06 |
| M203($\sim 5 \times 10^{-6}$ M) | 0 | 69.44 |
| | $\sim 0.5 \times 10^{-5}$ M | 55.75 |
| | $\sim 1.0 \times 10^{-5}$ M | 49.51 |
| | $\sim 1.5 \times 10^{-5}$ M | 30.06 |
| | $\sim 2.0 \times 10^{-5}$ M | 10.34 |
| M210($\sim 5 \times 10^{-6}$ M) | 0 | 79.10 |
| | $\sim 0.5 \times 10^{-5}$ M | 76.75 |
| | $\sim 1.0 \times 10^{-5}$ M | 53.31 |
| | $\sim 1.5 \times 10^{-5}$ M | 40.72 |
| | $\sim 2.0 \times 10^{-5}$ M | 33.68 |

### Raman spectrum of **β**-lactamaseproteins

Raman intensity of wild type and mutant β-lactamases (5µM) were measured in presence of ascending concentration of CAZ (5 µM-15 µM). Results showed a gradual increase of Raman intensity of β-lactamases with the increase of the concentration of CAZ in comparison with the wild type protein. There are three regions (880 cm^-1^, the region of Tyr doublet due to H-Bonding),(1360 cm^-1^, CH2- CH2 deformation) and (1640-1680 cm^-1^, characteristics of amide peak) in the spectrum which indicated the secondary structure alternation due to molecular interaction. A small blue shift was also observed for the mutant β lactamases in comparison with the wild type due to secondary structure adjustment during interaction with CAZ. (**Fig. S5**)

### Energy transfer between CAZ and β lactamases

The absorbance spectrum of CAZ (excited at 267 nm) and fluorescence spectrum of the TEM 1 β- lactamase at 340 nm overlapped with each other which satisfied the requisite condition of FRET between the CAZ and the proteins **(Fig. S6)**. Moreover from the Förster theory of energy transfer, the molecular interactions between the drug-protein complexes were evidenced from discrete values of the energy transfer efficiency (E); 0.47, 0.94, 0.79 and 0.82 for CAZ-wild type, CAZ-M184, CAZ-M203 and CAZ-M210 respectively which facilitated binding of CAZ to the mutants than the wild type.

### In silico study

Earlier *in silico* study from our laboratory indicated effective binding of CAZ to the wild type and mutant β lactamases (Mukherjee S K et al, 2018). The stability and conformational changes of the β lactamase in presence of CAZ were analyzed by MD simulation study.The movement of atoms and molecules for 100 ns of simulation were monitored. In presence of CAZ, the wild type β lactamases (1ZG4) and M184 exhibited C-α backbone deviations at about ∼1.5 to 1.75 Å and ∼1.75 to 2.5 Å respectively throughout the simulation trajectory and an equilibrium was maintained after 40 ns for both. However, for M203 and M210 C-α backbone deviations from ∼1.5 to 2.5 and 1.75 to 2.5 Å with an elevation at 60 ns for the latter which reached equilibrium from 70 ns was observed. The C-α all residue deviations for M184 and M210 were ∼2.5 to 3.25 Å and ∼2.5 to 3.40 Å respectively with an elevation at 60 ns and equilibrium from 70ns in case of the latter. In contrast, M203 showed reduced Cα backbone deviation at 30 ns akin to wild type throughout the 100 ns simulation trajectory. However, M203 showed C-α all residue deviations from ∼2.25 to 3.0 in comparison to wild type in presence of CAZ (**Fig. 5 A-F**)

**Fig. 5.**
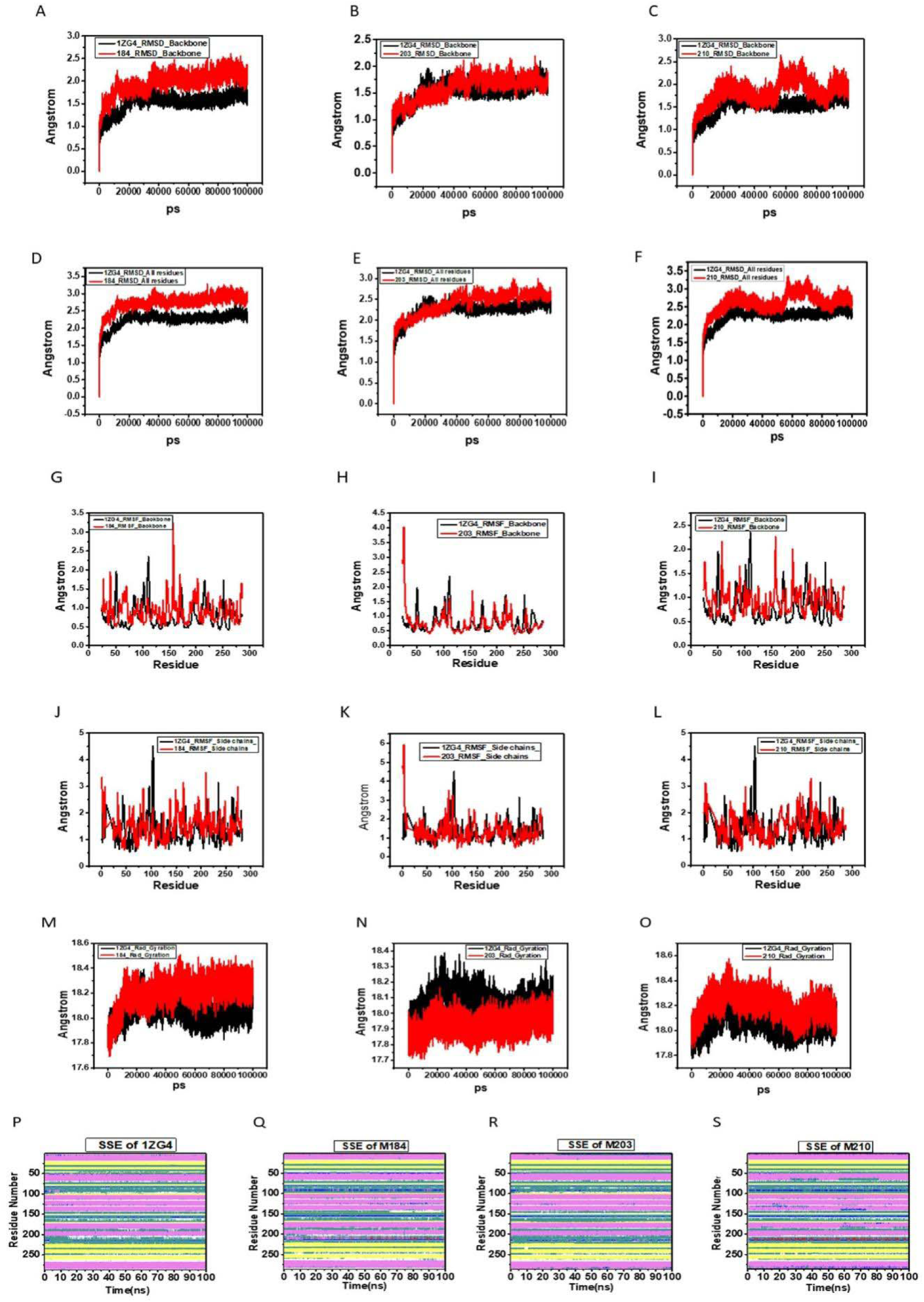
Cα backbone RMSD, Cα all residue RMSD, RMSF for Cα backbone, RMSF for Cα side chain and Rg for backbone Cα for 100 ns simulation time frame shown for the wild type TEM-1 (1ZG4) interacting with CAZ; displayed in black, in comparison with mutant proteins M184 (A,D,G,J,M), M203 (B,E,H,K,N), and M210 (C,F,I,L,O); displayed in red respectively. Projection of SSEs throughout the MD simulation trajectory in presence of CAZ for wild type TEM-1 (IZG4) (P), M184 (Q), M203 (R), M210 (S). SSE composition is represented by different color codes: helices; pink, turns; green, 3-10 helix; blue, β-strands; yellow, coils; white.

The residue level fluctuations (RMSF) of Cα backbone for M184 and M210 in presence of CAZ were moderately high (∼3.25 Å) in comparison to wild type at residues located between 50-80, 120-170, 200, and 225-250 respectively. However, a comparable pattern of fluctuation was observed for M203 and wild type except at residue 150. Similarly, RMSF for Cα side chain of the mutant proteins (M184, M203, M210) in presence of CAZ compared to wild type in presence of the drug showed similar pattern as the RMSF for Cα backbone (**Fig. 5 G-L**).

The compactness of protein-drug interaction was ascertained by Cα Radius of gyration (Rg) plot. The residue level Rg value fluctuation for wild type protein was within the range of ∼17.8 to 18.3 Å, whereas for mutant protein (M184) was within the range of ∼17.8 to 18.2 Å and reached equilibrium from 40 ns onward in presence of CAZ. Furthermore, in presence of the drug the Rg value fluctuations for M203 was ∼17.8 to 18.1 across the 100 ns simulation trajectory but for M210, the Rg value fluctuations was ∼17.9 to 18.6 Å with a decrease in fluctuation within the range of ∼18.2 to 18.4 Å from 60 ns onward to attain the equilibrium. The overall Rg analysis for compactness in the mutants in presence of CAZ showed lowest radius of gyration that demonstrated tight binding of CAZ to the mutant proteins in comparison to wild type (**Fig. 5 M-O**).

The protein-drug interactions were further monitored by estimation of secondary structure elements (SSE) composition (helices, strands, turns, loops) in the interacting proteins across the simulation trajectory throughout the 100 ns time frame. SSE for each amino acid residue of the mutants exhibited significant structural adjustment in comparison with wild type in presence of CAZ. In M184 increase of 310-helix in residue positions 100, 160, 220, turns in residue position 50, 90, 160, coils in position 140, 200 and π-helix in positions 210 to 225 was evidenced. In M203, the segment of the residues in positions 60-90 showed increase of 310-helix, coil structure at residues 150-170 and 225-240. M210 showed an increase of α-helix and coil with the absence of π-helix at positions 60-90 and 160-170 in comparison with the segment of residues in the wild-type with increase of 310-helix and increase of π-helix at residue position 210-225 respectively (**Fig. 5 P-T**).

## 5. Discussion

Emergence of varied β-lactamases was reportedwhich was the primary cause of β-lactam resistance in uropathogenic *E. coli*, the primary etiologic agent of urinary tract infection. These β-lactamases were found to harbor mutations which paved the possibility of effective binding of the different β-lactams and their inhibitor combinations to develop resistance. Therefore, studies on the protein-drug interactions have become increasingly important from a pharmaceutical perspective.Absorption spectroscopy study provide information on the binding modes of ligands with proteins pave the possibility of ground state complex formation between protein-ligand interfaces (Swain et al, 2020). The individual UV-Vis absorption spectrum of the CAZ and TEM-1 β-lactamase were markedly different to that of CAZ-β-lactamase complex, indicating the formation of new complexes between CAZ and β-lactamases. Appearance of isosbestic points at 339.4 nm, 360.2 nm, 341.4 nm and 340 nm for the TEM-1-CAZ, M184-CAZ, M203-CAZ and M210-CAZ systems respectively suggested that an equilibrium was established between the respective two species and reflected the formation of complexes between them. Therefore, these observations signify that the ground state complexes are formed between respective wild type and mutant β-lactamase proteins and CAZ. The UV-Vis absorption of the four β-lactamases increased and a slight red-shift that occurred with an increase in concentration of CAZ, which indicated that, the drug-protein interaction resulted in subtle conformational changes in the protein in presence of the drug. A hyper- chromic effect and slight red-shift of the maximum absorption peak were also separately observed with increasing CAZ concentrations in solution. The shift of maximum wavelength signifies hydrophobic effect on Trp residue due to complex formation between drug and respective proteins (Topala et al. 2014). However, the observations obtained from absorption studies were not sufficient to study the interactions in detail.

Therefore, other excited state spectroscopic techniques e.g., fluorescence spectroscopy, TCSPC etc. were preferred to study the binding mode of protein-drug interactions. Variation in temperature produced a local change in the protein microenvironment as heat disrupted the hydrogen bonds, non-polar hydrophobic interactions, which might produce a local change in the Trp microenvironment and affected the spectroscopic properties of the protein. In this study, the increase of quantum yieldof wild type and mutant β-lactamases with the ascending temperatures represented the change in the local hydrophobicity in presence of CAZ. Moreover, a decrease in fluorescence intensity (quenching) of the β-lactamases (wild type and mutants) with increase in temperature in presence of CAZ alsoindicated formation of a new complex.which corroborated with the change in the protein microenvironment (Topala et al. 2014,Ghisaidoobe et al.2014,Butler et al.2015.Bardhan et al. 2011)

Additionally, in CAZ-β lactamases system the quenching mechanism might also be either due to simultaneous occurrence of static (ground state) and dynamic (excited state) interaction modes. The static quenching had already been established by ground state complex formation. However, the decrease in fluorescence lifetime of wild type and mutant β lactamases with simultaneous addition of CAZ accounted only for dynamic quenching supported by the linear Stern-Volmer plot. Moreover, the positive deviation of Stern-Volmer plot for CAZ-β-lactamase systems demonstrated the simultaneous occurrence of static and dynamic quenching. Furthermore, the Stern-Volmer plots of the wild type β-lactamase in presence of increasing concentration of CAZ at different temperatures indicated a gradual shift towards dynamic mode but for mutant-CAZ complexes a sharp shift towards dynamicity indicated a more rigid interaction of the drug to the mutant enzymes (Mukherjee S K et al, 2021Bardhan et al. 2011)

Determination of binding constant and binding stoichiometry of β-lactamases in presence of CAZprovided evidence for both static and dynamic quenching.The thermodynamic parameters of binding, i.e., changes in standard enthalpy (ΔH^0^), entropy (ΔS^0^) and Gibb’s free energy (ΔG^0^) using van’t Hoff isotherm exhibited the nature of interacting forces between drug and protein. Usually, four types of forces play an important role in drug–protein interaction, like electrostatic forces, hydrophobic forces, van der Waals interactions and hydrogen bonding (Ross et al.1981). In this study it was found that the interaction of CAZ with the mutant proteins was predominantly hydrophobic, however van der Waal’s interactions and hydrogen bonding was the primary interacting forces in the binding of CAZ to the wild type enzyme. Furthermore, the anisotropy changes demonstrated flexible interactions of the drug with the mutants than the wild type protein which also supported predominance of hydrophobic interaction between the drug and mutant enzymes (Zhao et al. 2019).

The circular dichroism (CD) is a sensitive technique to monitor the conformational changes in proteins upon interaction with ligand molecules (Bertucci et al.2010). CD spectra of the wild type and mutant proteins in presence of CAZ further corroborated conformational changes in the secondary as well as tertiary structure alterations in the protein at pH 7.0. Additionally, assessment of energy transfer efficiency values (E) using FRET technique and Raman spectroscopy provided evidence that the specific mutations in the M184, M203 and M210 assisted an effective binding of CAZ to the mutant β lactamases than to the wild type protein (Chakraborty et al 2015,Lee et al 2016,,Pezzotti et al 2021).

*In silico* MD simulation analysis of the wild type and the mutant β lactamases revealed increase in the RMSD for the C-α backbone for the mutant β lactamases which might be attributed to the overall conformational flexibility of the protein to accommodate the drug molecule (Mukherjee et al.,2021,Gill et al. 2019]. Therefore, the overall comparison of RMSD plots for wild type and the variants showed that the C-α backbone residues in the latter maintained stabilizing interaction with the drug molecule.

Similarly, RMSF for Cα side chain of the mutant proteins (M184, M203, M210) compared to wild type in presence of the drug showed similar pattern as the RMSF for Cα backbone which further documented the enhanced conformational flexibility in the mutant β-lactamases to bind to CAZ than the wild type (Mukherjee et al.,2018,Fantini et al 2020). The Cα Radius of gyration (Rg) for compactness in the mutantβ lactamases in presence of CAZ also demonstrated a tight binding of the drug to the mutant proteins in comparison to wild type (Mukherjee et al.,2021,Galdadas et al.2021). Furthermore, the residue-based conformational changes in the mutants in presence of CAZ also indicated that the altered secondary structure elements (SSE) in the mutants probably imparted additional overall stability and rendered conformational flexibility compared to the wild type protein (Fantini et al 2020). This *in-silico* study established that the mutant-CAZ interactions were more stable and rigid with minimum conformational alterations between the interacting residues compared to the wild type-CAZ interaction (Yang et al 2020).

## Conclusion

This is the first of its kind research that illustrated the comprehensive bioactive interaction of CAZ with three novel mutants of TEM β-lactamases derived from clinical isolates of uropathogenic Escherichia coli at the molecular level via spectroscopic and biophysical approach. β-lactamases remain at the forefront of driving antibiotic resistance via hydrolysis of β-lactam antibiotics, necessitating the understanding of their evolving mechanisms. We explained the role of far-site mutations in clinical TEM β-lactamase variants that enhance ceftazidime (CAZ) binding and resistance, even though they are distant from the active site. Through absorbance and fluorescence spectroscopy, coupled with biophysical measurements, we demonstrated that the mutations favor tighter, mostly hydrophobic interactions with CAZ, as opposed to hydrogen-bond-dominated binding in the wild-type enzyme. In silico structural modeling also revealed that the mutations confer conformational flexibility, enabling more rigid and stable accommodation of CAZ within the enzyme’s active site. These findings underscore the way that non-active site mutations can indirectly contribute to antibiotic resistance by optimizing substrate binding dynamics. The widespread therapeutic application of β-lactams likely selects and propagates such variants since they provide a survival advantage under therapeutic pressures. This study highlights the difficulty of resistance evolution and the need for new strategies to inhibit mutational adaptations outside of traditional active-site-directed drug design.

## Significance

This study represents a pivotal advancement in understanding the molecular evolution of antibiotic resistance by describing how distant mutations in TEM β-lactamase variants enhance ceftazidime (CAZ) resistance, a critical third-generation cephalosporin. Through spectroscopic and biophysical approaches, this study describes how these mutations facilitate tighter, hydrophobic-restricted binding to CAZ compared to hydrogen-bond-dependent binding of the wild-type enzyme which opposes the traditional paradigm that resistance results from active-site modifications alone.The findings emphasize an important evolutionary strategy that, mutations far from the catalytic site can modulate the enzyme architecture to enhance substrate specificity.This study highlights the urgent necessity for new approaches to fight against resistance mechanisms beyond active-site targeting. By integrating structural, functional, and evolutionary data, the study provides a road map for designing next-generation antimicrobials to combat the relentless rise of resistance in Gram-negative bacteria.

