## Supplemental File for "Distant Site Mutations in Clinical TEM Beta-Lactamase Variants Enhance Non-Covalent Binding to Ceftazidime: Insights from Spectroscopic and Biophysical Investigations"

### Supplementary data

Fig. S1. Steady state UV vis absorption spectra of (1) CAZ (20 µM ), (2) TEM β (5 µM)µM), (3) summation of the spectra of (TEM + CAZ), (4) difference between the spectra of mixture of (TEM and CAZ) and sum of (TEM and CAZ and (5) spectra corresponding to the mixture of TEM with CAZ.

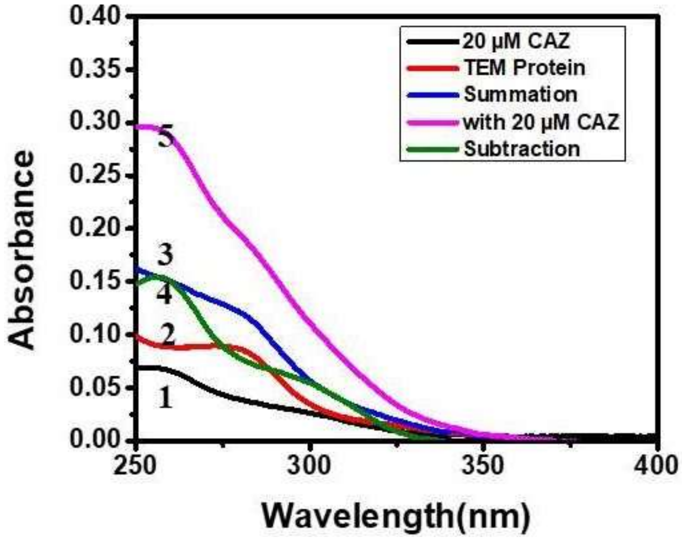

Fig S2: Thermal denaturation curve wild (A) TEM-1, and mutant (B) pUE184TEM, (C) pUE203TEM, (D) pUE210TEM β-lactamases at different temperature (298K, 303K.308K and 311K) with 0.5 mM concentration.

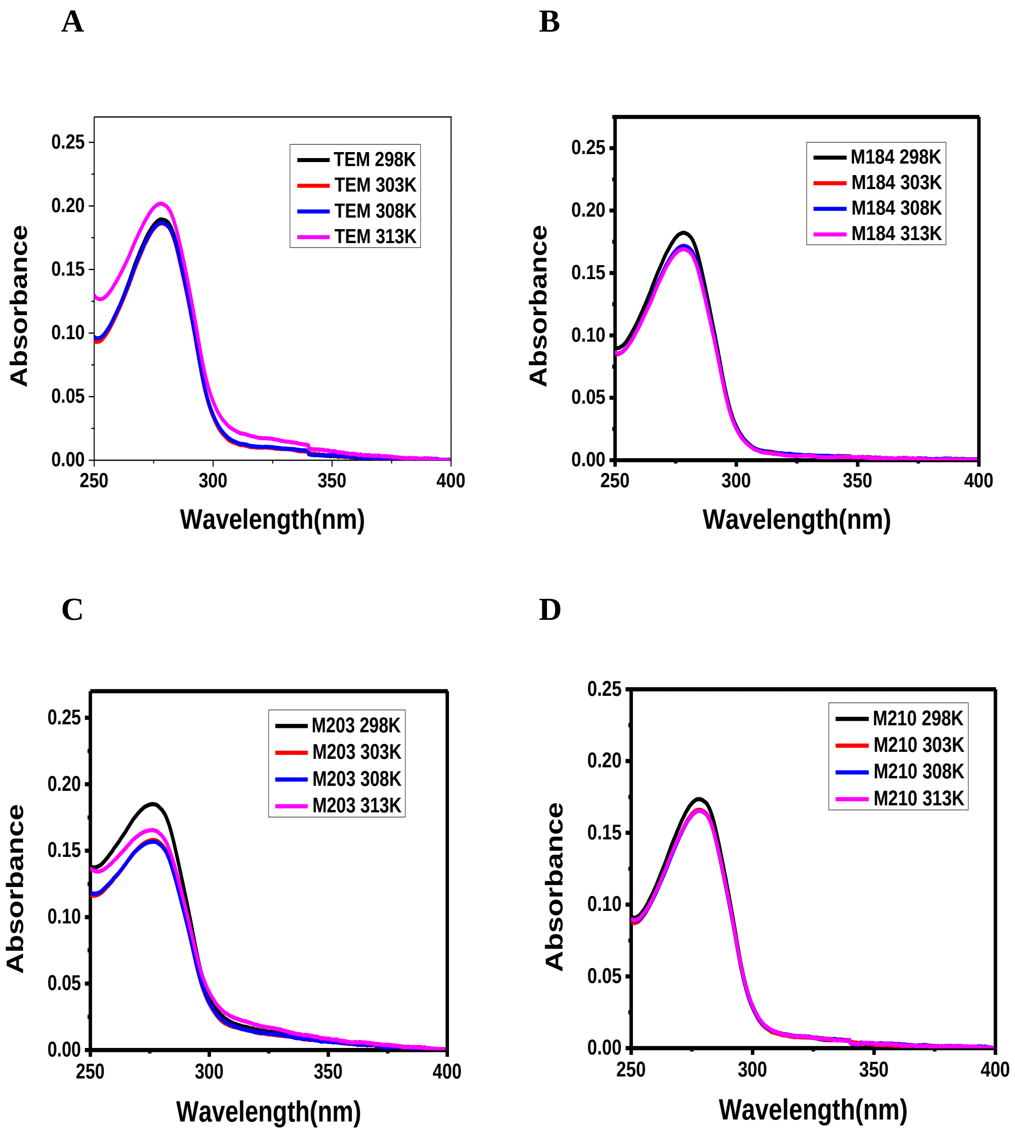

Fig. S3. Fluorescence excitation and emission spectra of **A,B** TEM-β lactamase (wild type) and mutant TEM β-lactamases **C,D** M184, **E,F** M203 and **G,H** M210 at increasing concentrations (5-20µM) of β-lactamases in presence of 5 µM concentration of CAZ at temperature 298K.

A

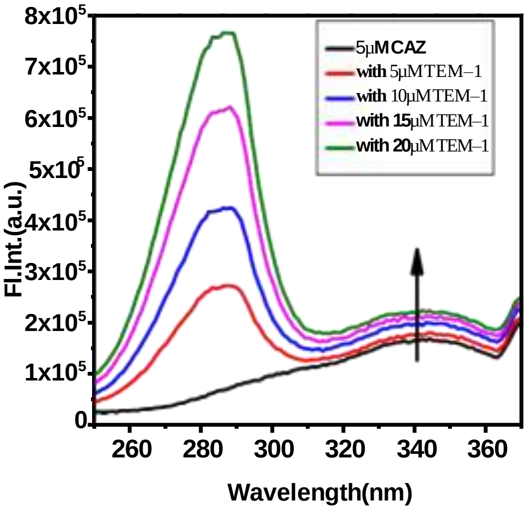


B

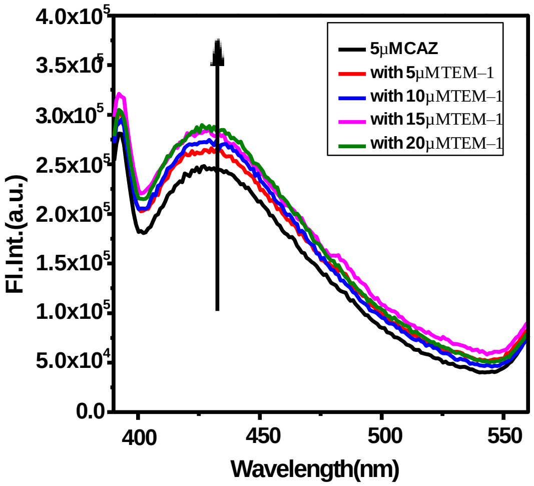


C

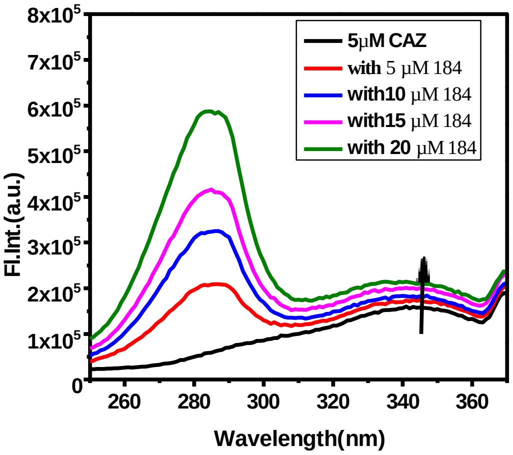


D

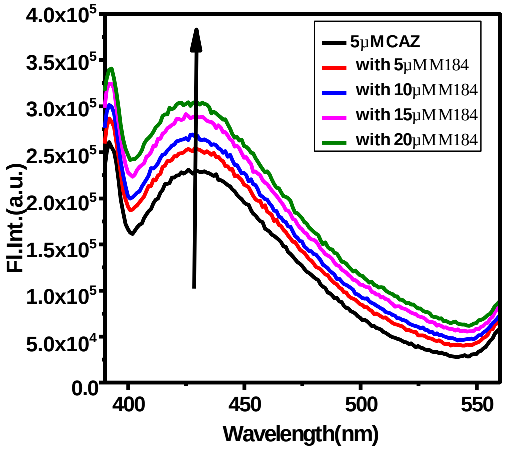

E

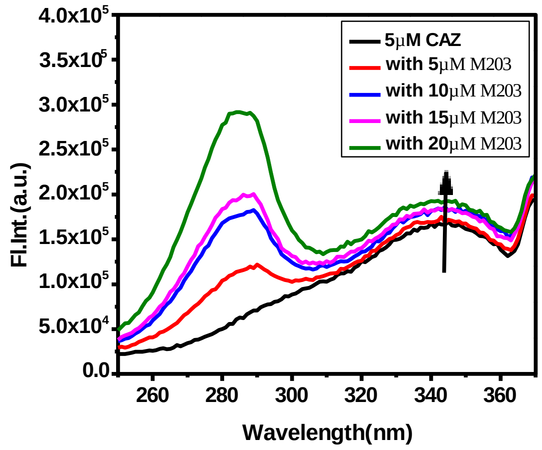

5μM CAZ
with 5μM M203
with 10μM M203
with 15μM M203
with 20μM M203
Fl.Int.(a.u.)
Wavelength(nm)


F

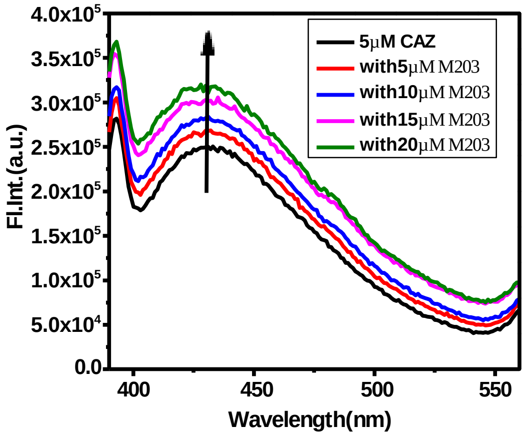

5μM CAZ
with5μM M203
with10μM M203
with15μM M203
with20μM M203
Fl.Int.(a.u.)
Wavelength(nm)


G

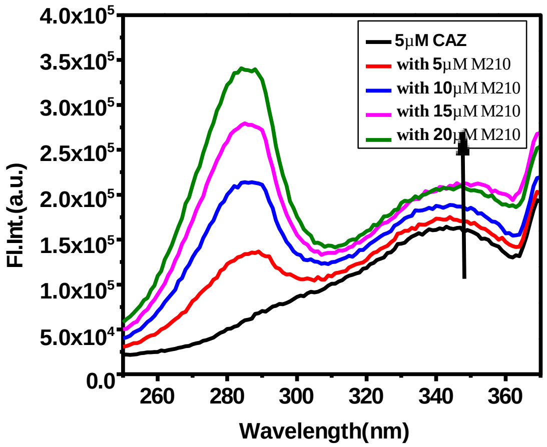

5μM CAZ
with 5μM M210
with 10μM M210
with 15μM M210
with 20μM M210
Fl.Int.(a.u.)
Wavelength(nm)


H

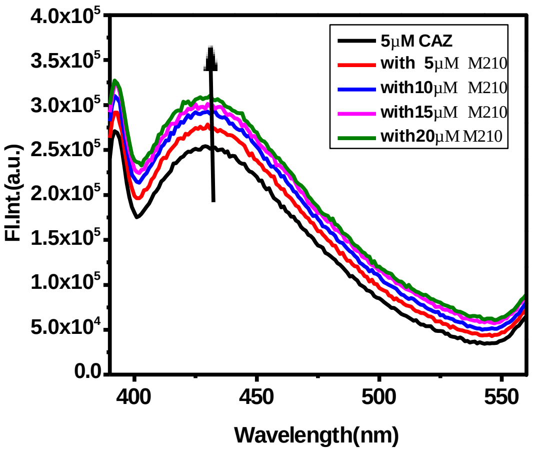

5μM CAZ
with 5μM M210
with10μM M210
with15μM M210
with20μM M210
Fl.Int.(a.u.)
Wavelength(nm)

Fig. S4. Far-UV spectra and near-UV spectra of **A, B** TEM β lactamase (wild type protein) and mutant TEM β lactamases **C,D** M184, **E,F** M203 and **G,H** M210 at 5 μM concentration in the presence of CAZ (0-20 μM) in sodium phosphate buffer (pH 7.4).

A

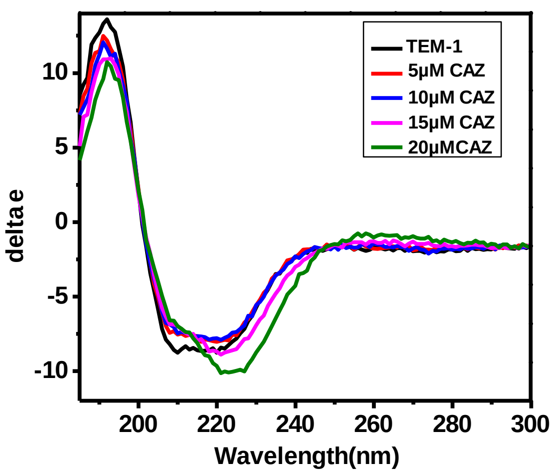


B

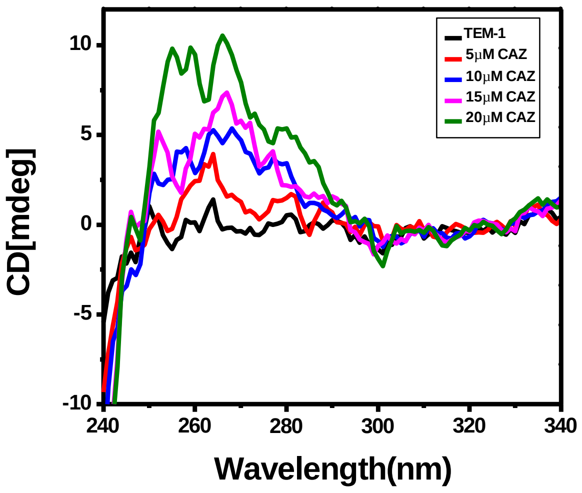


C

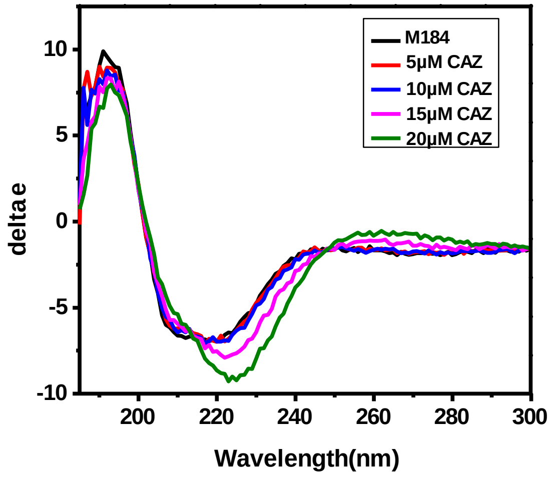


D

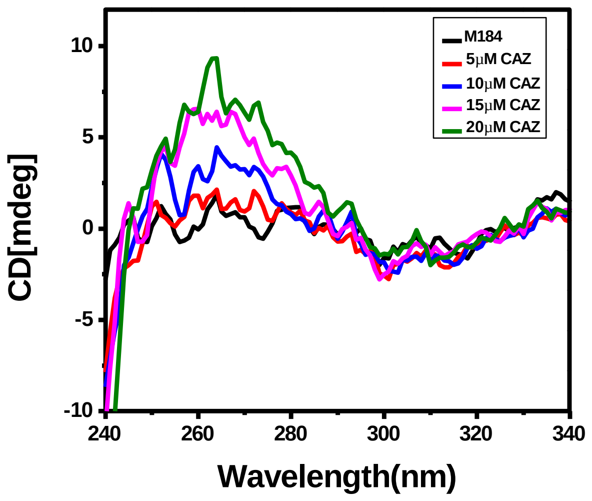

E

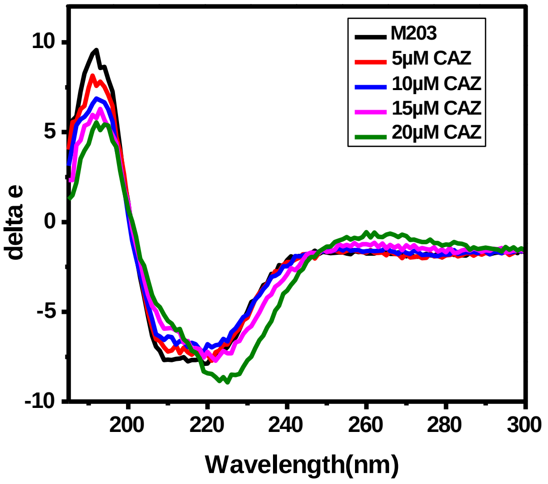
M203
5μM CAZ
10μM CAZ
15μM CAZ
20μM CAZ
delta e
10
5
0
-5
-10
200
220
240
260
280
300
Wavelength(nm)

F

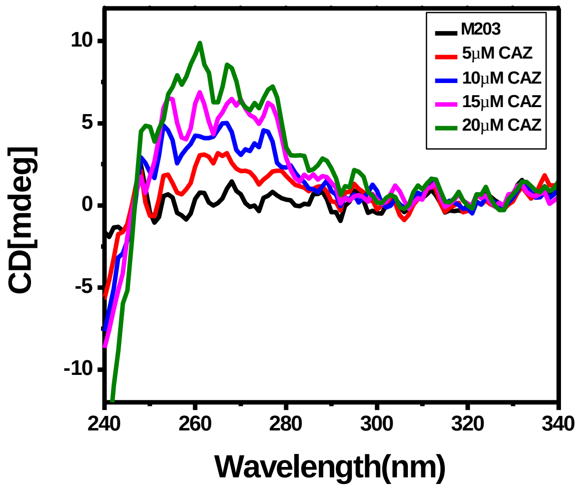
M203
5μM CAZ
10μM CAZ
15μM CAZ
20μM CAZ
CD[mdeg]
10
5
0
-5
-10
240
260
280
300
320
340
Wavelength(nm)

G

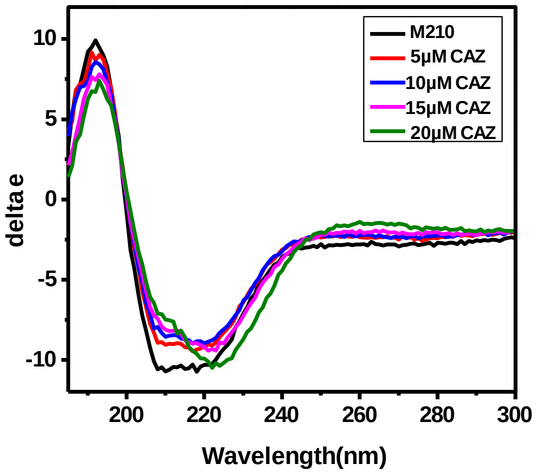
M210
5μM CAZ
10μM CAZ
15μM CAZ
20μM CAZ
delta e
10
5
0
-5
-10
200
220
240
260
280
300
Wavelength(nm)

H

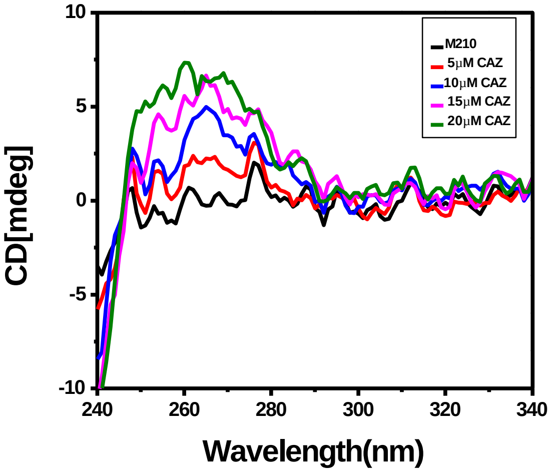
M210
5μM CAZ
10μM CAZ
15μM CAZ
20μM CAZ
CD[mdeg]
10
5
0
-5
-10
240
260
280
300
320
340
Wavelength(nm)

Fig. S5. Representative Raman spectra of **A**TEM-β lactamase (wild type protein, 1ZG4) and mutant TEM β-lactamases **B** M184, **C** M203, **D** M210 at 5 µM in presence of increasing concentration of CAZ (0-15 µM) at temperature 298K

A

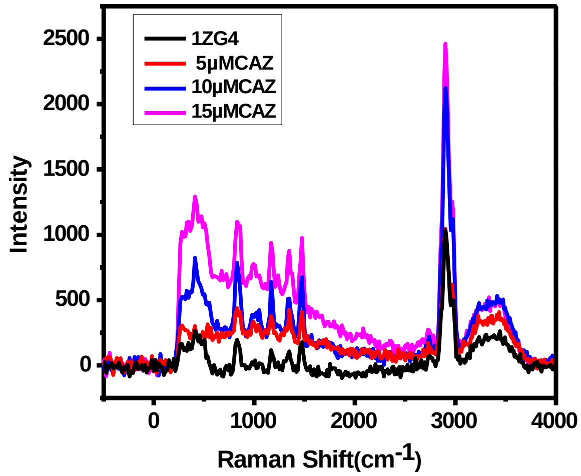


B

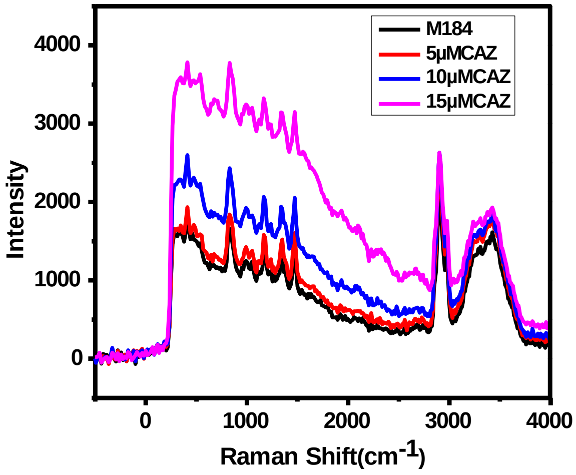


C

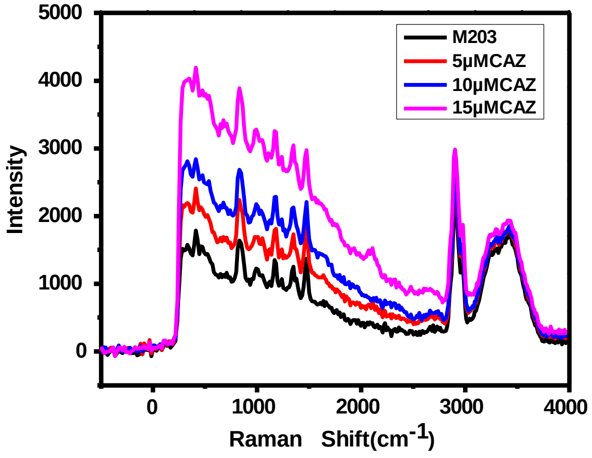


D

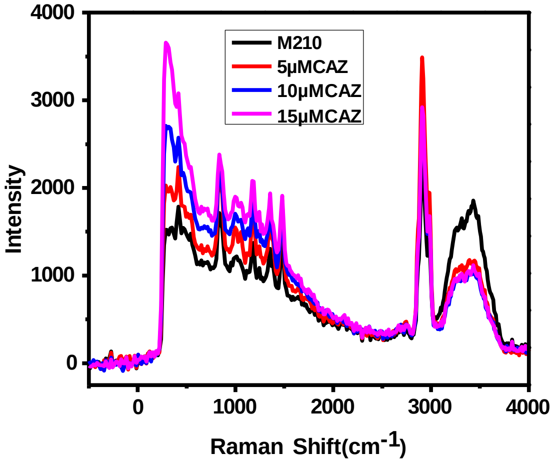

Fig. S6. Overlapping absorption spectrum of CAZ (acceptor) and the fluorescence spectrum of TEM-1 β-lactamase(donor)

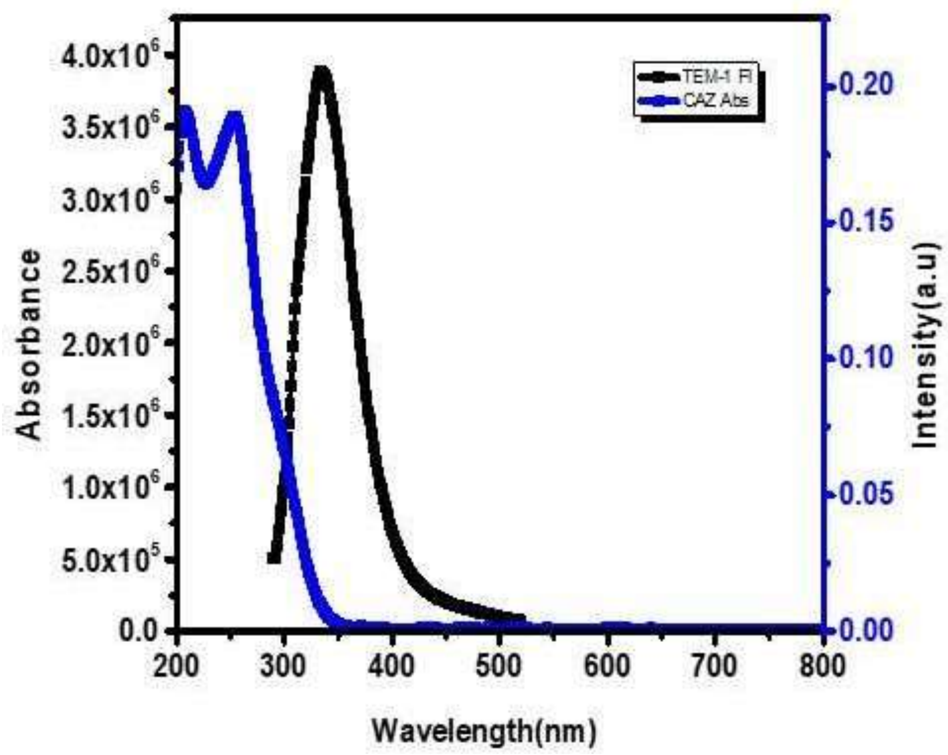
